# In vivo Quantification of Neural Criticality and Complexity in Mouse Cortex and Striatum in a Model of Cocaine Abstinence

**DOI:** 10.1101/2022.08.02.501652

**Authors:** Wesley C. Smith

## Abstract

Self-organized criticality is a hallmark of complex dynamic systems at phase transitions. Systems that operate at or near criticality have large-scale fluctuations or “avalanches”, the frequency and duration power of which are best fit with a power law revealing them to be scale-free and fractal, and such power laws are ubiquitous. It is an attractive concept in neuroscience since spiking avalanches are exhibited by neural tissue, and may underpin how minuscule events could scale up to circuits and provide adaptive psychobiological function. Much is yet to be understood about criticality *in vivo* in the healthy brain and in disorders such as addiction, as drugs may alter the critical state’s “tuning” to generate drug seeking and dysphoria. Thus, here a novel toolset was developed to use neural avalanches and their self-similarity, rather than power law fit slope exponents as is canonically done, to quantify criticality in a previously collected high-density electrophysiological *in vivo* corticostriatal dataset from a mouse model of early cocaine abstinence. During behavioral quiescence, in the prefrontal cortex but not ventral striatum of cocaine-dosed mice, it was found that critical tuning is enhanced compared to drug-free controls. Additionally, an empirical biological demonstration of complexity’s theoretical correlation to criticality was shown in drug-free mice, was exponentially enhanced in drug-treated cortex, but was absent in the drug-treated striatum. As shown, quantifying criticality grants experimental support for the “critical brain hypothesis” and allows for statistical interpretation of inter-subject variability and development of further testable hypotheses in systems neuroscience.

**Significance Statement:** The “critical brain hypothesis” asserts neural networks are comparable to material in phase transitions at a critical point, their “avalanches” of system-wide spike bursts best seen in log-log plots of probability vs. avalanche size or duration, with slope following a scale-free or fractal power law. In discussing criticality, “critical tuning” is mentioned but quantification thereof left for later experimentation, despite being necessary for a scientific hypothesis. Presented are methods to quantify critical tuning through assessing similarity or fractalness among corticostriatal avalanches collected using high-density electrophysiology in cocaine-conditioned mice, along with an empirical *in vivo* confirmation of the mathematical concept that data complexity correlates with criticality. Interestingly, cocaine enhances critical tuning in cortex and aberrantly modifies complexity in a region-specific manner.

## Introduction

An emergent property of clusters of neurons, self-organized criticality (SOC) may be the most fundamental expression of a neural network responding to perturbations in a scale-free manner (1, 2), allowing for high dynamic range in sensory responses and increased complexity in information encoding (3, 4); neural ensembles may benefit from this property and guide adaptive or maladaptive behaviors (5–7). Being ubiquitous, SOC potentially is the framework uniting descriptions of various interacting self-similar natural systems like earthquakes, wildfires, birds flocking, and ferromagnets (8–10). Emerging evidence, perhaps unsurprisingly, suggests the brain’s information structure obeys related principles of statistical mechanics (11), an extension of thermodynamics.

Although consilience of evidence supports criticality in physics and chemistry, as an explanation for neuroscientific problems it remains to be tested rigorously (3, 4, 12–15). Briefly, SOC systems have scale-free fluctuations, “avalanches”, which irrespective of size or duration, are statistically distributed following a power law seen in log-log coordinates (16–18). Neural avalanches are identified with a characteristic recording array-wide spiking profile, bounded by brief, typically 5ms, periods of inactivity. Power laws fits have the same slope at any two points and are thus scale-free, ergo the deformation of the system through avalanches should follow a fractal structure, as in cracking noise (8, 19).

The last two decades of brain criticality experimentation in preparations as disparate as murine tissue culture, *in vivo* primate physiology, and human EEG and fMRI have shown consistent functional aspects of criticality (17, 20–23). The “tuning” of neural slices towards and away from criticality via pharmacological manipulations, inferred by the changing of avalanche distribution from exponential to a power law, has revealed maximum information transfer and dynamic range of response to weak inputs at the critical point (24–26), and other experiments have shown a similar principle *in vivo* in rats that awaken from anesthesia (27). Human imaging data corroborates this and has shown loss of critical dynamics concomitant with propofol-induced loss of consciousness (28). More recent murine work *in vivo* follows a similar path in discussing critical tuning in terms log-log fit exponents (29), consistent with the usage of Hurst exponents and detrended fluctuation analysis (DFA) to ascertain if criticality has a functional role in physiological neuroplasticity and in disease states (15).

Although criticality is thought to be an integral part of consciousness, sensory response, and attention (30), there is nothing known about how addictive drugs like cocaine alter criticality despite it being well-accepted that systemically taken narcotics change qualia through psychosis (31–33), craving contributes to dysphoria (34), and withdrawal has vast central and autonomic consequences (35), with addiction having correlates at synaptic and circuit levels (36). Additionally, there is no study demonstrating criticality in the striatum, a centrally important subregion in dopamine-dependent learning and memory, despite the depth of electrophysiology regarding it (37, 38); neural criticality could provide an aid in understanding systems correlates of decision making and addiction-dependent errors.

Most work to date assessing critical tuning is limited to essentially fits of power laws (13, 39) or correlations in ensemble fluctuations, which can provide conflicting results (40, 41) since neurons have inconsistent firing and weak correlations (42). Fit exponents are tangential to criticality because observing ubiquitous power laws doesn’t prove system criticality since even successive fractionation or even noise can yield power law distributions, and there’s no consensus of sufficient conditions for ascribing criticality to the brain (43). To address these shortcomings, alternative post-hoc analysis was performed on corticostriatal 512-channel awake-behaving *in vivo* silicon probe electrophysiology datasets in mice undergoing brief cocaine abstinence after behavioral conditioning (44), using only epochs of behavioral quiescence prior to cue exposure.

To compare results between canonical methods and the current high-density *in vivo* approach, basic avalanche properties are first described similarly to established criteria (45) in both regions. Then, to quantify tuning, a proxy metric for self-similarity and fractalness in avalanches, the median absolute error (MAE) of all mean avalanche collapse shapes per subject, rather than an arbitrary subset (14, 29, 45) is proposed. Additionally, correlation matrices of collapse shapes corroborate MAE as valid metric to assess how self-similar scale-free avalanches are. It was hypothesized that cortical avalanches in the drug treated group are more fractal (with a lower MAE) as the slight increase in quiescent firing previously seen in cocaine-dosed cortices (44), rather than hypoactivity, may signal ensemble bursting being enhanced, and thus altered avalanches and criticality being enhanced. This was validated; however, striatum appeared to be less modulated by cocaine in terms of criticality and complexity. Since a rigorous quantification of spike-time shuffling’s effects on avalanche structure was unavailable (14, 29, 45), applying 1ms of Gaussian jitter destroyed avalanche structure in nearly all animals, further evidence of the exquisite temporal structure of neural avalanches. Finally, by quantifying criticality, it was amenable to mathematical correlation to complexity, and shown that in controls there was a roughly linear correlation between MAE and complexity in cortex and striatum, which after drugs was aberrantly altered in a region-dependent manner. Overall, presented is further empirical support for deep brain *in vivo* neural criticality, an additional method to quantify oft-overlooked inter-subject variability in addiction (44, 46, 47), and a method to enable further hypothesis testing in neural criticality.

## Results

### Awake in vivo recording paradigm and demonstration of avalanche shapes in vivo

Figure 1A is a schematic of the recording; for detailed explanations of conditioning, see reference (44). All subsequent analysis refers to the behavioral quiescence period of 15 minutes prior to any cue onset, as discussed in that paper. All recordings utilized a 512-channel 3D silicone probe array inserted into the prefrontal cortex (Fig. 1B, left) and ventral striatum (Fig. 1B, right) at the indicated anterior-posterior coordinates. These probe arrays yield extensive firing rasters (Fig. 1C), with two smaller avalanches (orange highlight for Fig. 1D & blue highlight for Fig. 1E) shown. Avalanches are defined as array-wide bursts bounded by 5 ms frames of inactivity; consistent with other groups, even these small avalanches have a characteristic inverted-U shape (45).

**Figure 1.**
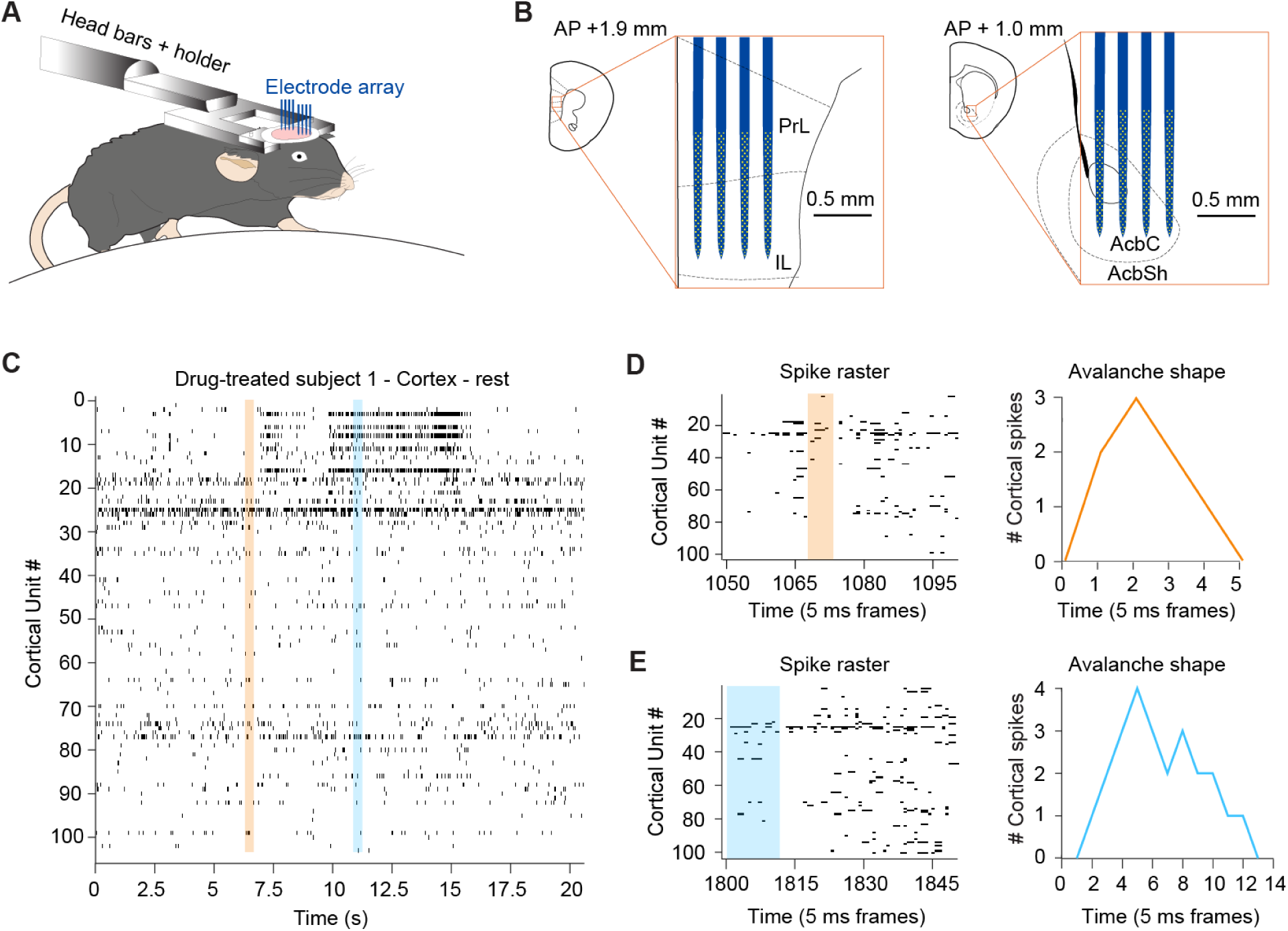
Awake in vivo recording paradigm and demonstration of avalanche shapes in vivo. (A) Head-fixed mice on a uniaxial treadmill were acutely recorded from, as previously in reference (44), and analyses performed on 15 minutes of data collected during quiescence. (B) Schematic of electrode array positioning in the cortex (left) and striatum (right), with scale and anterior-posterior coordinates indicated (PrL = Prelimbic cortex, IL = Infralimbic cortex, AcbC = Accumbens Core, AcbSh = Accumbens Shell). (C) Segment of a single animal’s cortical firing raster. (D) Expanded raster of the orange trace in (C) (left) and the corresponding avalanche shape (right); time is indicated in 5 ms frame bins. (E) Expanded raster of the blue trace in (C) (left) and the corresponding avalanche shape (right).

### Scale-free properties of in vivo corticostriatal avalanche data

Avalanches in each region had a duration (frame count) and event power or magnitude (spikes per frame) assigned, and these were plotted using a MATLAB histogram and logfit function; Fig. 2A shows representative striatal avalanche size data, while 2B shows corresponding cortical data. As indicated in both panels, the goodness of log-log fit R^2^ was above 0.85 in both brain regions, with avalanche event power slope “τ”∼-3/2, near the golden ratio, theoretical calculations, and neurobiological observations of this value (3, 45, 48, 49). For the same subject, duration “α” and power vs. duration “Γ”, the latter also known as “α-1/τ-1” or “β” (29, 45, 49), plots are in 2C and 2D, respectively. Drug-exposed and control τ values did not differ in a statistically significant way for the cortex (Fig. 2E, left; Mann–Whitney U test, p = 0.3518) or striatum (Fig. 2E, right; Mann–Whitney U test, p = 0.8522), but are within physiological parameters of SOC systems, evidence that this custom toolbox can reliably extract acute avalanche data from silicon probe datasets (50). Perhaps most importantly, the mathematically derived and empirically observed Γ values are also near canonical observations in critical systems and similar amongst each other (Kruskal-Wallis test, p = 0.7129), further evidence towards critical tuning (49) and the validity of the mathematical relationships of α, τ, and Γ in these data. Despite not finding differences in log-log fit exponents between drug treatments, systems operating at criticality have more features such as fractal fluctuations in their data, and many systems that do not operate at criticality exhibit α similar to critical data (51), demonstrating that fit exponents are not sufficient to demonstrate criticality, and *a priori* may not provide enough information for meaningful quantitative testing. However, these results do suggest the data follow a power law and may be fractal.

**Figure 2.**
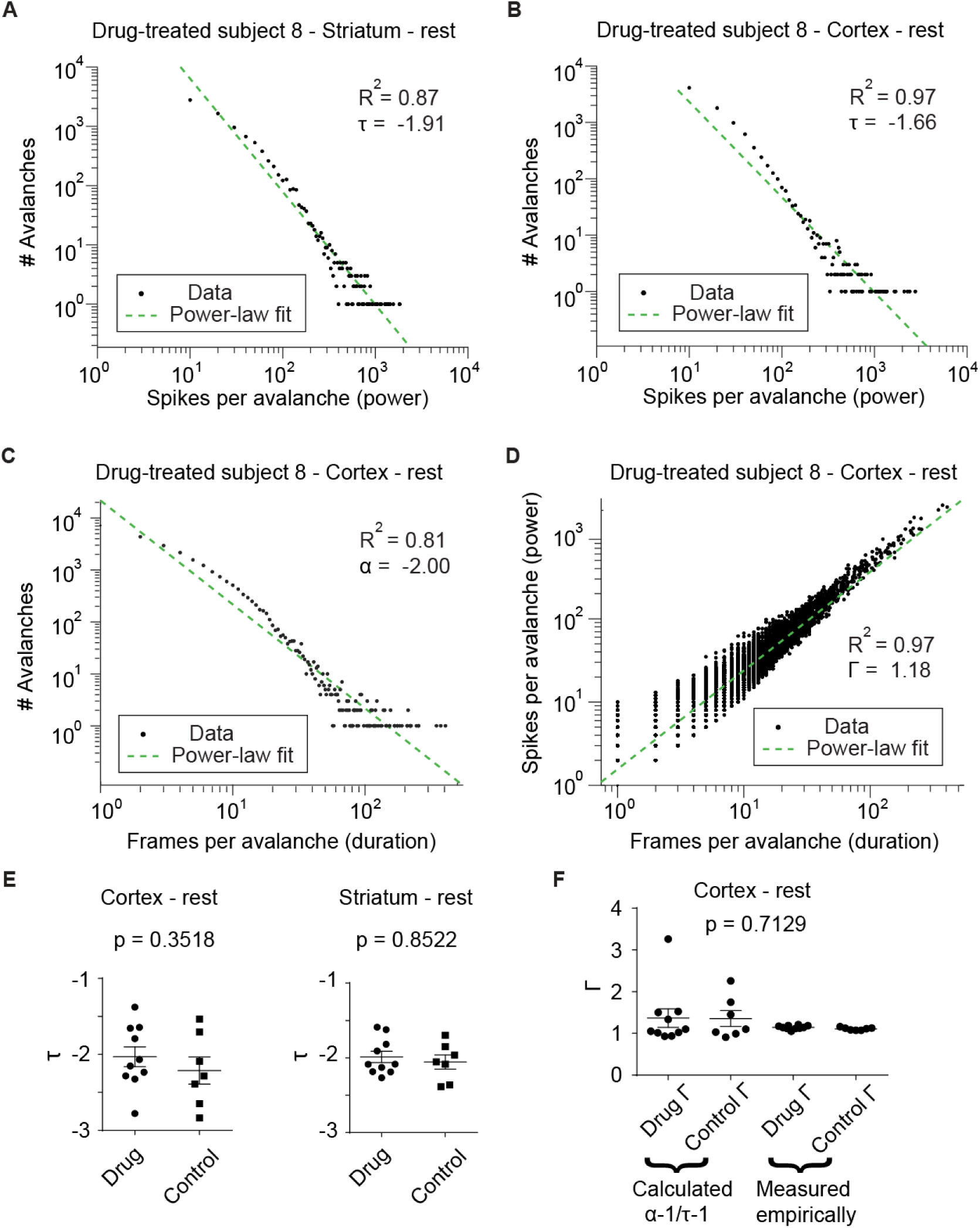
Scale-free properties of in vivo corticostriatal avalanche data. (A) One subject’s striatal avalanches plotted with avalanche count vs. event power in log-log coordinates follow a power law with R^2^ = 0.87 and τ = -1.91. (B) Cortical avalanches plotted as in (A) with R^2^= 0.97 and τ = -1.61; both sets of values have high goodness-of-fit and τ consistent with other scale-free critical phenomena in nature and neurophysiological data. (C) As in (B), but showing avalanche duration follows a power law with R^2^= 0.97 and α = -2.00, while (D) shows a logfit plot of power vs. duration from (B) and (C), with R^2^= 0.97 and measured Γ = 1.18; panels A-D are all based off one subject’s data. These indicate *in vivo* data are highly consistent with previous observations of critical phenomena obeying power laws. (E) Resting-state τ did not differ between drug (n = 10) and control group (n = 7) in cortex (left, Mann–Whitney U test, p = 0.3518) nor striatum (right, Mann–Whitney U test, p = 0.8522), while in (F) calculated and measured cortical Γ are not statistically different in any condition in the cortex (Kruskal-Wallis test, p = 0.7129), but do show all exponents are mathematically consistent with each other.

### Avalanche collapses and median absolute error (MAE) to quantify critical tuning shows enhancement in cortex after cocaine treatment and requires sub-millisecond spike coordination

To better assess critical tuning, a toolbox was made to take all avalanches and collect those that had the same duration and collapse them using fractional duration and z-scores for spiking. If the data were fractal and critically tuned, then avalanches of any shape length should look the same, and the level of similarity or dissimilarity can be quantified. Figure 3A demonstrates six mean avalanche shapes of different durations, but it should be noted that there are many dozens of avalanche duration groups per animal and brain region even in a 15 minute recording (see Fig. 2C). One animal’s entire avalanche criticality profile at rest can be visualized by then obtaining the mean of all normalized mean shapes and plotting the standard error; **Supplementary** figure 1 **contains** a stepwise schematic of this process. Each regional dataset can be treated similarly, and one can roughly see a qualitative difference in normalized avalanche collapse profiles in drug-treated mice (Fig. 3B) and control mice (Fig. 3C).

**Figure 3.**
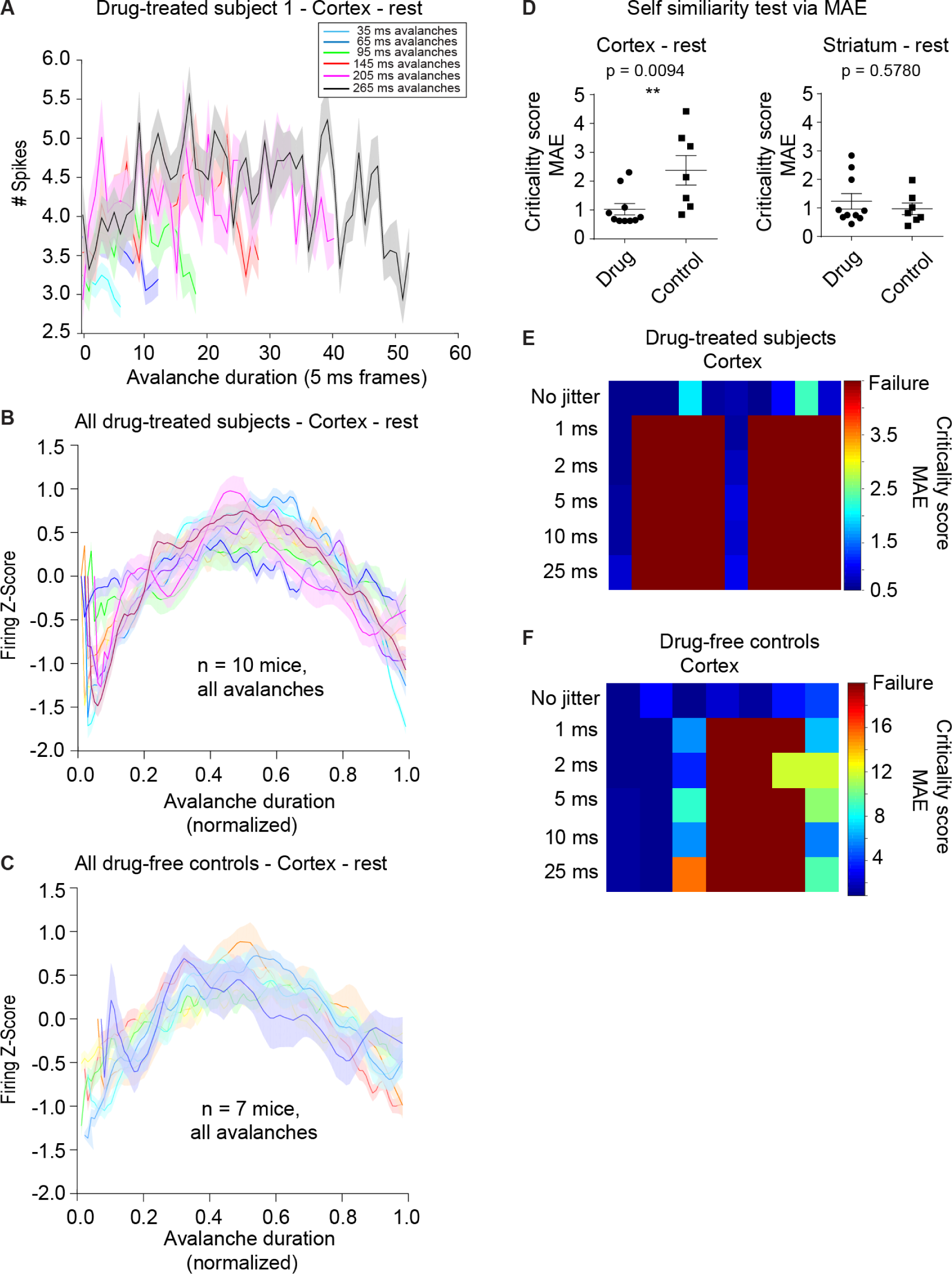
Avalanche collapses and median absolute error (MAE) to quantify critical tuning shows enhancement in cortex after cocaine treatment and requires sub-millisecond spike coordination. (A) Six mean avalanche shapes are plotted (shaded areas: SEM), comprised of suprathreshold –avalanches of indicated duration. Generally, long and short avalanches appear similar. (B) Normalized shape profile for cortex in drug-treated mice. (C) Normalized shape profile for cortex in controls. (D) Cortical data at rest have a lower MAE and are more tuned towards criticality in the cocaine-dosed mice (left, Mann–Whitney U test, p = 0.0094) whereas there is no similar effect observed in the striatum (right, Mann–Whitney U test, p = 0.5780). (E) As little as 1 ms of Gaussian jitter on spiking data in drug-treated mice can cause the algorithm to fail to detect avalanches in 8/10 animals. (F) The same jittering method fails to detect avalanches or markedly shifts MAE to non-physiological levels in 5/7 controls.

Two quantitative tests on these data were performed to ascertain differences between resting-state critical tuning in drug-treated and control mice in cortex and striatum, which is the chief advantage of this toolbox. The first test relies on median absolute error (MAE), one of a number of summary statistics that are used to measure data dispersion (52), to see how critically tuned each brain region in each animal was. MAE can be written as:

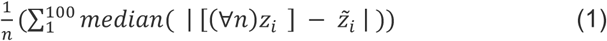

Where n is the number of collapse shapes in that condition, [(∀*n*)] refers to a matrix of all n z-scores, *z*∼ is the median z-score, and *i* is the time within the nth normalized, interpolated collapse shape, which ranges from 1-100. Since this is a unitless number representing absolute error, the smaller it gets for each animal in each condition, the more similar each median collapse shape is to each other, and thus, more self-similar or fractal.

Using this MAE metric, it was found that in the cortex, during behavioral quiescence and in absence of cocaine, mice that had experienced drug conditioning were more tuned to criticality than drug-naïve controls (Fig. 3D, left; Mann–Whitney U test, p = 0.0094); this did not hold true in the striatum (Fig. 3D, right; Mann–Whitney U test, p = 0.5780). This is not intuitive, as one familiar with addiction and cognitive bias regarding addicts (53) would expect hypoactivity (46) and less computationally beneficial brain states in cocaine users absent drug. However, the current result is in line with slight quiescent cortical hyperactivity (44) and data on recently abstinent cocaine users having compensatory cortical activity (54) prior to the transition to hypoactivity (55).

One might object that observing and quantifying avalanches is based on a coincidence without functional relevance, or is at most a biophysical curiosity. To respond to this potential objection, other groups have demonstrated that large amounts of Gaussian jitter applied to spiking data disrupt any avalanches (14, 45), but used jitter as large as 5 ms, an order of magnitude larger than most pyramidal neuron spike widths (56). Here it is shown that as little as 1 ms of Gaussian jitter can cause the algorithm to fail to observe any avalanches in the cortex of drug-treated mice (Fig. 3E) and in controls (Fig. 3F) in nearly all mice, and in those few cases where avalanches were still observed, an increase in MAE was concomitant with the jitter applied, consistent with previous work (45). Thus, the functional relevance of avalanche fractal structure and sub millisecond-level timing in “signal-less” quiescent recordings 15 min long (57) is indeed curious, at the very least.

### Correlation matrices of mean interpolated collapse shapes complementarily show cocaine enhances critical tuning in the cortex

To answer another potential objection to usage of a proxy for avalanche fractalness, in each animal pairwise correlations for each interpolated collapse were performed, such that low duration collapses would be compared to longer and longer collapses, and vice versa. The resulting pairwise correlation matrices are based on control data (Fig. 4A; only cortical data shown, Pearson’s R shown in top right, p-values of each pairwise shown in bottom left of each plot) and drug group data (Fig. 4B; each plot constructed identically to 4A), per animal. This is mathematically one of the simplest ways to demonstrate similarity in timeseries data, but comes with the caveat of interpreting the deluge of correlation coefficients.

**Figure 4.**
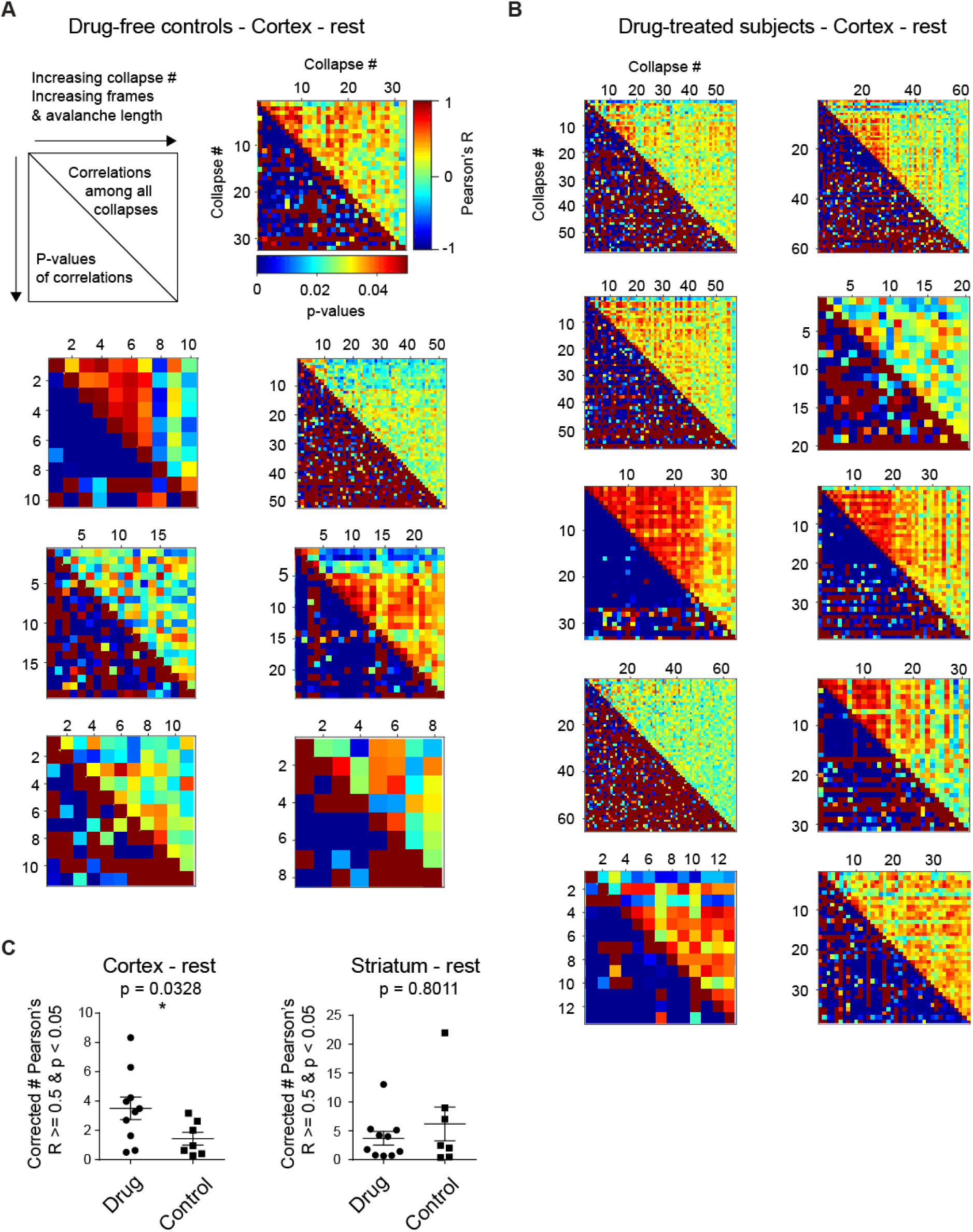
Correlation matrices of mean interpolated collapse shapes complementarily show cocaine enhances critical tuning in the cortex. (A) Control cortical data plotted as interpolated collapse correlations and a legend for interpreting each dataset (top left). Note that the cutoff for significance is p = 0.05, and in most cases, p-values were very low (p < 0.01) or above 0.05. (B) Each drug-treated subject’s cortical interpolated collapse correlation matrices with the same legend as in (A). (C) Cortical data at rest have a higher count of statistically significant positive correlations > 0.5 in the cocaine-dosed mice (left, Mann–Whitney U test, p = 0.0328), whereas striatal data show no such difference (right, Mann–Whitney U test, p = 0.8011).

Subsequently, counts were made of all statistically significant (p < 0.05) Pearson’s R values > 0.50 in the drug treated vs. control condition, and a statistically significant increase in these counts for drug treated mice was found in cortex (Fig. 4C, left; Mann-Whitney U test, p = 0.0328) and again, not striatum (Fig. 4C, right; Mann-Whitney U test, p = 0.8011). These data are consistent with the MAE result, showing that even when directly comparing collapse shapes to each other, it is found more often that collapse shapes are correlated, and thus, self-similar and closer to being fractal in the drug-treated condition. These data also suggest if any malleability in the system occurs early in transition to addiction, the first place it could manifest is the cortex.

### Cocaine increases spiking complexity in the cortex and aberrantly alters the physiological correlation of complexity to criticality in a region-dependent manner

As outlined above, criticality and information entropy and system complexity are related. By comparing total individual neural entropies to the joint entropy of the system, the degree of coordination, dubbed integration of neural firing can be measured (58), which is similar to the dynamic correlation or “total correlation” that DFA approximates in systems thought to be operating at criticality. A publicly available complexity toolbox was used (13) on these current resting state data with the hypothesis that in the cortex of cocaine-treated mice, complexity would be enhanced concomitant with MAE, and that in the physiological condition absent drug, complexity should be correlated with criticality in both brain regions, with this experiment offering the opportunity to test mathematic theory *in vivo*.

Complexity represented by integration is naturally difficult to interpret, so a demonstration of complexity is presented in a 12 pseudo-neuron system with three model constraints: Random spiking (Fig. 5A, top), ordered spiking (Fig. 5A, middle), and complex spiking (Fig. 5A, bottom). The resulting integration of the random data reveals no coordination, similar to white noise (Fig. 5B, top), while the ordered state has a perfectly linear integration curve (Fig. 5B, middle), indicating that although there are interactions, there is no variability or entropy, sometimes referred to as “surprise” (59), in them. Only complex data have a nonlinear integration (Fig. 5B, bottom), suggesting variable coordination and nonzero complexity.

**Figure 5.**
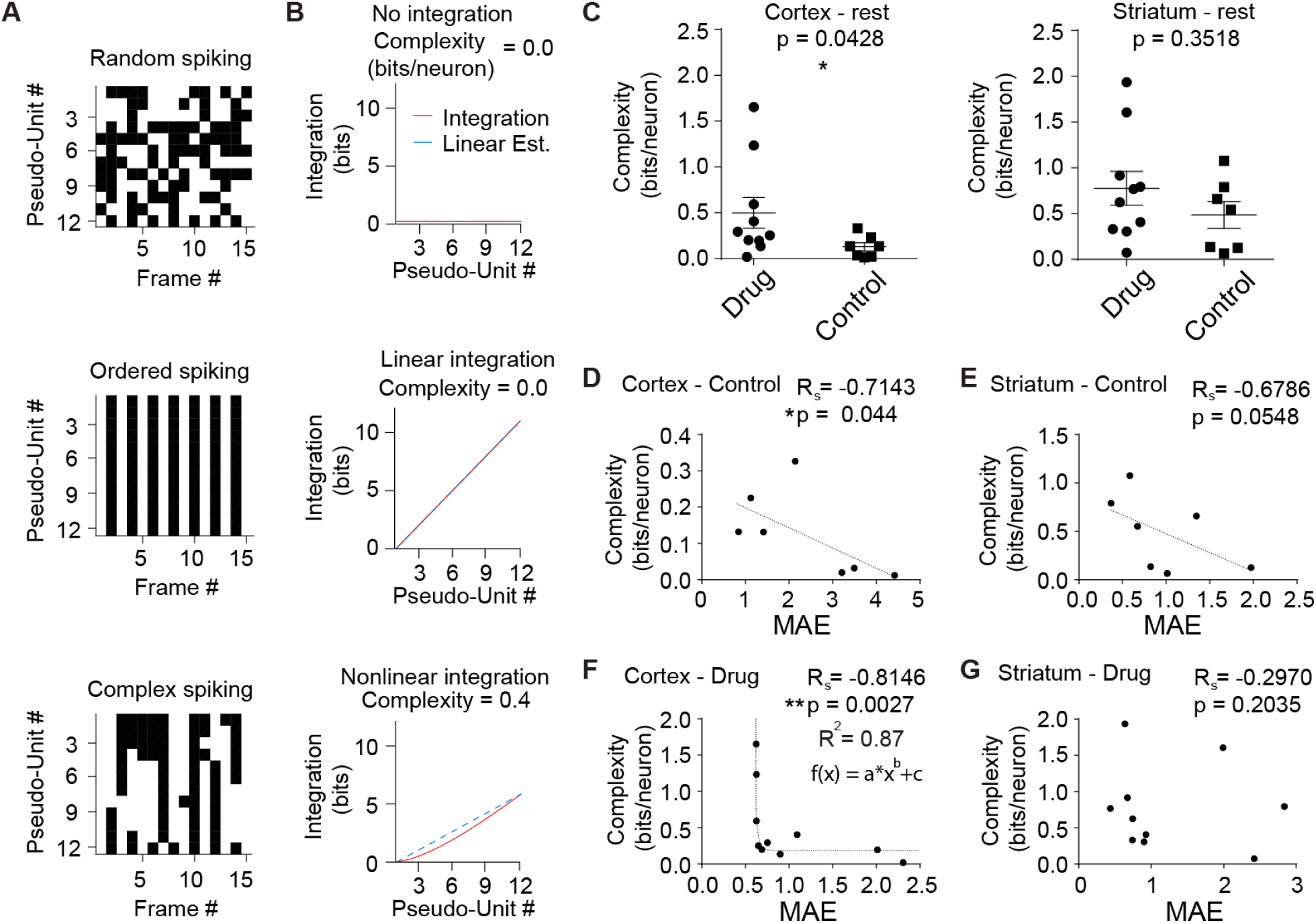
Cocaine increases spiking complexity in the cortex and aberrantly alters the physiological correlation of complexity to criticality in a region-dependent manner. (A) Complexity example with pseudodata spike rasters under three different regimes: Random (top), ordered (middle), and complex (bottom). (B) Integration plots and complexity values based on the pseudodata in (A). Complex data (bottom) have a nonlinear, scale-variant integration, while random data (top) have no integration, and ordered data (middle) have perfectly predictable integration. Thus, only complex data have nonzero complexity. (C) Cortical data at rest have a higher complexity score in the cocaine-dosed mice (left, Mann–Whitney U test, p = 0.0428), while striatal data lack a statistically significant difference between drug and control groups (right, Mann–Whitney U test, p = 0.3518). (D) In control mice, complexity was inversely correlated to MAE in the cortex (R_s_ = -0.7143, p = 0.044), meaning that as data have more critical tuning with a lower MAE, complexity increases. (E) Control striatal complexity also trended towards inversely correlating with MAE (R_s_ = -0.6786, p = 0.0548). (F) Correlation between MAE and complexity was aberrantly enhanced in the cortex of drug-treated mice (R_s_ = -0.8146, p = 0.0027) and changes from being best fit by linear approximation to an exponential as indicated. (G) No statistically significant correlation between MAE and complexity was observed striatum of drug-treated mice (R_s_ = -0.2970, p = 0.2035), indicating cocaine treatment aberrantly disrupts the correlation in (E).

Using this method it was found that drugged mice have enhanced complexity in the cortex, compared to control mice (Fig. 5C, left; Mann-Whitney U test, p = 0.0428), but not in the striatum (Fig. 5C, right; Mann-Whitney U test, p = 0.3518). Additionally, as expected, a strong correlation was found between MAE and complexity in these resting-state data in the cortex of controls (Fig. 5D; R_s_ = -0.7143, p = 0.044) and to a lesser extent the striatum (Fig. 5E; R_s_ = -0.6786, p = 0.0548). Surprisingly, after cocaine exposure, the correlation was exponentially enhanced in cortex (Fig. 5F; R_s_ = -0.8146, p = 0.0027; R^2^ = 0.87, fit using MATLAB’s “*curvefit*” where f(x) = a*x^b^ +c) and broken in the striatum (Fig. 5G; R_s_ = -0.2970, p = 0.2035). As this is the only result seen that runs counter to expectations about drugs and criticality, it could be that this is one of the features that dictates aberrant drug induced plasticity at the systems level during early abstinence.

## Discussion

In trying to explain a curious result about cortical hyperactivity rather than hypoactivity after cocaine conditioning (44), a novel method to detect neural avalanches and quantify critical tuning was developed, issues regarding defining critical tuning in the “critical brain hypothesis” were addressed (4, 43, 51), and evidence for critical tuning in deep brain structures was presented alongside documentation of effects of a potent narcotic on neural criticality and complexity. This work represents the first usage of nanofabricated multi-shank electrode recordings to characterize deep brain criticality analysis *in vivo* in awake, behaving mice. Even the most advanced *in vivo* murine recordings used in avalanche analysis only had 32 channels with a 200 μm inter electrode spacing and 400 μm inter-shank distance (27), with the recorded sites in superficial cortex in anesthetized subjects. Other recent work used avalanche frames that are an order of magnitude greater with an order of magnitude fewer electrodes (29). Comparatively, the inter-electrode spacing here is almost an order of magnitude closer, with an order of magnitude more electrodes on shanks half the distance apart. Despite these distinctions, avalanches were reliably extracted using criteria set by other groups, lending credence to the concept of SOC as a ubiquitous feature of neural systems.

When plotting avalanche size in log-log coordinates, power law fits similar to others’ were replicated, with similar τ and α exponents near the golden ratio (3) and corresponding physiological critical Γ exponents. Interestingly, despite having dosed mice with cocaine over a week, fit exponents appeared equivalent to control mice, which might be due to the differences in recording technology as outlined above, or in how the 5 ms inter-avalanche interval was designated. Others have used an interval dependent on a subject’s neural activity levels (60), which may more accurately delineate avalanches. Additionally, multiple other potential model fits and advanced log-fitting tools specifically optimized to these data weren’t employed, thus fits may deviate from what other groups might obtain using more advanced mathematics (11, 61). On the other hand, using a stock log-log fit toolkit to obtain exponents with R^2^ > 0.85 implies success. Ultimately, power laws are only one facet of the neural criticality hypothesis, and whether they vary due to drug treatment does not falsify the fractal results examined. As it currently stands, few studies on pharmacological modification of critical tuning exist, and have only worked with bath applications of channel blockers in organotypic slices, for example (24, 25). It would make sense that τ would be altered in those experiments, and in light of these studies, the current result is less surprising, given that incubation of craving is much more physiologically subtle and physiological mechanisms exist *in vivo* that could be compensatory.

The quantification of interpolated mean avalanche collapses allows for testing that lend support to the “critical brain hypothesis”, as previous discussions of critical tuning relied on shape fitting or methods tangential to how fractal the collected data are, since fit exponents are uninformative and occasionally spurious. Importantly, when using MAE or correlation matrices of collapses, data are not transformed beyond normalizing, and both directly quantify dissimilarity or similarity, respectively. The metrics used here are well understood, computationally simple, and open up data to further easily interpretable analyses.

The observation of quiescent enhancement in criticality through use of two mathematically different metrics only in the cortex of drug-treated mice isn’t intuitive at first glance, given the abundance of data supporting cortical hypoactivity (55) and the stereotype of an addict is one who is aimless, lacking in self-control, or in withdrawal and unable to think clearly. A parsimonious explanation for this result is that only during the transition to addiction there is an increase in quiescent firing; following this, hypoactivity disengages the cortex, and in the literature there exists support for this scenario in recently abstinent subjects (54). That the striatum’s critical tuning is unaffected is somewhat expected, given that previously no resting-state firing changes were observed in it following this protocol (44). Maladaptive plasticity within the striatum during incubation is thought to take place over multiple weeks (62), and thus how incubating craving affects criticality remains an open question.

Consistent with criticality being enhanced at rest in the cortex after brief cocaine abstinence, a measure of complexity was also enhanced in the same group of mice, and as was recently hypothesized, degree of critical tuning and complexity are correlated in both the cortex and striatum of controls (14). Notably, this correlation breaks down in striatum in the cocaine-dosed mice, but is greatly enhanced in the cortex. The aberrant interplay between complexity and criticality could be a system-state change that denotes a transition to addiction, concomitant with previous results in the same subjects revealing a spontaneous increase in corticostriatal LFP coherence between 25-45 Hz (44).

Ultimately, these results represent a great step closer in understanding inter-subject variability in how both addiction and neural criticality manifest *in vivo.* At its core, this work enhances the criticality hypothesis in deep brain structures with state-of-the-art high-density, awake *in vivo* data. Strong evidence is presented for early stage resting cortical hyperactivity after cocaine treatment, rather than hypoactivity, as well as multiple quantitative metrics to address avalanche collapse shape fractal qualities, rather than relying on power laws only. Jittering the data in a manner not done previously also demonstrates how critical timing is to the exquisite the structure of avalanches, and how unlikely the presented results are to be spurious, given how many time bins were taken and how many spikes observed. Finally, the empirical biological validation of complexity mathematics warrants further investigation in the SOC field.

## Materials and Methods

As these analyses are based on previous data (44), no animals were sacrificed for these post-hoc tests to be in best accordance with the “3 Rs” principle in animal research (63). 10 mice were part of the cocaine-treated group and 7 were part of the saline-only group, and all testing was in accordance with UCLA’s IACUC. All recordings were performed in the absence of drug or saline injections, 24 hr after the final conditioning session.

### Neural avalanche criteria and statistical testing of universal critical dynamics

Criteria established by Friedman, et al. were adapted to observe universal critical dynamics in cortical slice culture (45). Avalanches were defined as region-wide bouts of activity in 5 ms bins from single spikes up to hundreds of spikes, bounded by 5 ms of inactivity (Fig. 1C-E). Only clearly-defined projection or fast-spiking neurons in either region were chosen. All avalanches were collected but only quiescent period (15 min prior to cue onset as in (44)) avalanches were analyzed, since post-cue periods would need concatenation to permit an analyzable epoch, breaking avalanches and treating early cues similarly to late cues regarding composition and encoding. Unlike other groups (45), all avalanche durations more than 25 ms long (5 frames) that had more than 19 avalanches in them were used; this is a more strict data requirement for testing data fractal quality. Avalanches were plotted with *logfit* in MATLAB (64) to obtain fits and exponents α, τ, and Γ ( (Fig. 2). Exponents were subject to rank-based tests to assess difference in exponents between treatment groups; all statistical tests were rank-based, unless otherwise indicated, due to the sample size in each group and inability to match samples (10 drugged mice vs. 7 controls).

Per brain region, the mean of each accepted avalanche duration type with suprathreshold avalanche count was obtained (a mean shape) following z-score (firing) and fractional duration (frame count) normalization and plotted with SEM per time point as in Fig. 3B&C. Each animal’s set of mean shapes was then assessed for similarity to each other through interpolating all mean shapes at 100 points per shape in MATLAB. Per animal MAE of all interpolated collapse shapes was obtained as a final variable representing fractal fit or best estimate of self-similarity using formula [1]. As this value decreases, the error between avalanche mean collapse shapes decreases, and thus, indicates more self-similarity. These data were used in ranked statistical tests as before to observe differences among drug and control groups in both regions (Figs. 3-5), and correlations between groups were made with Spearman’s correlation (R_s_). To address additional mathematical concerns regarding repurposing MAE for self-similarity, correlation matrices of collapse shapes for each animal were made (only quiescent cortex shown in Figs. 4A & 4B) and counts of all statistically significant (p < 0.05) pairwise Pearson’s R values > 0.50 were compared between the drug-treated group vs. the control group as a broad method of examining self-similarity (Fig. 4C); note that these counts were corrected for the count of collapse durations included in each animal to not overweigh animals that had more viable collapses. Finally, a system-state entropy-based test for complexity (13) in the firing patterns of quiescent period data in each animal, per region, was performed and the resulting metric used in a comparison between treatment groups and correlated to MAE (Fig. 5)

### Data availability statement

All data and custom code are available from the author upon request.

## Acknowledgments and funding sources

The author would like to thank S. Masmanidis, N. Timme, M Porter, J.Y Kim, and I.J. Cho for support and discussions regarding this work, and the Brain Science Institute in The Korea Institute of Science and Technology for providing additional feedback. Portions of this manuscript were presented as part of the author’s doctoral dissertation, to which the author retains copyright. Data collection and initial analysis for this research was funded by the National Institutes of Health T32DA024635 predoctoral training grant on drugs of abuse, the Alfred P. Sloan Foundation, the McKnight Technical Innovations in Neuroscience Award, National Science Foundation Chemical, Bioengineering, Environmental, and Transport Systems Grant 1263785, and National Institutes of Health R01 DA034178. The current analyses and presentation of this work was supported by the Republic of Korea Ministry of Science Information and Communications Technology /National Research Foundation Brain Pool Fellow Program Grant #2021H1D3A2A02082997 and The Korea Institute of Science and Technology.

---

**Supplementary Figure 1.**
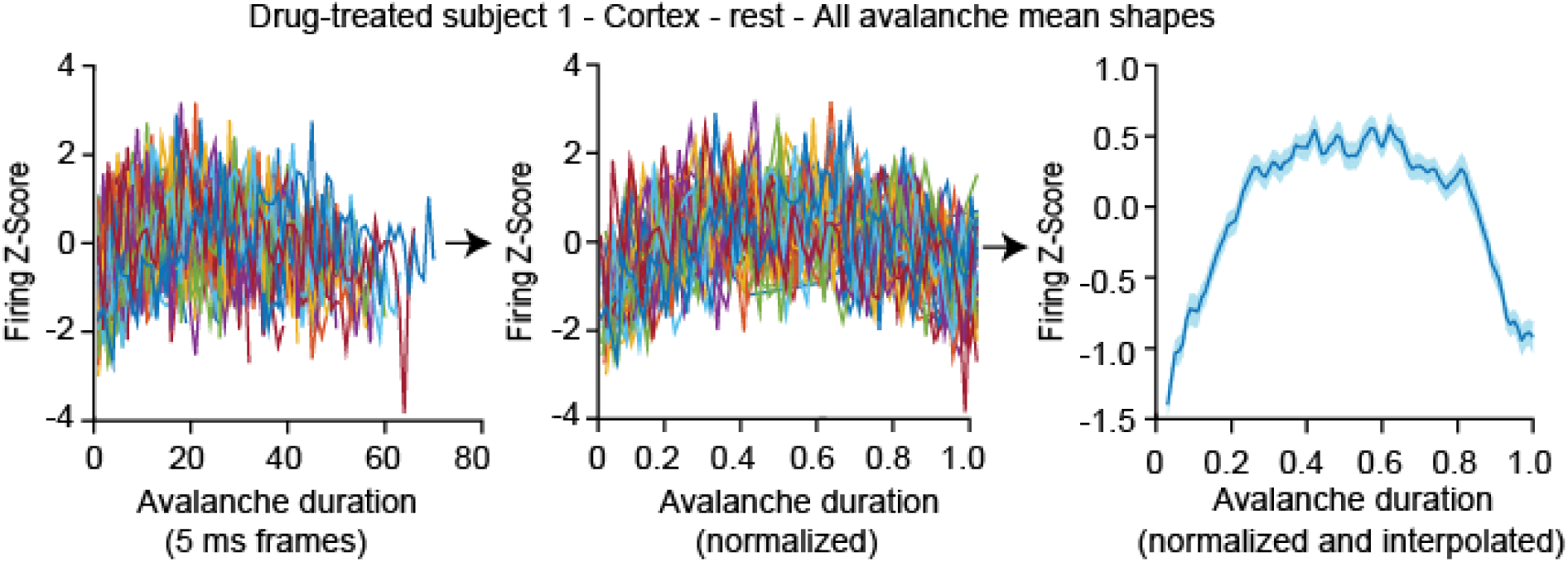
Schematic of how each subject’s avalanche profile is obtained. The collapse process starts by taking all mean avalanche shapes (finding all viable avalanches of differing durations that meet threshold requirements and normalizing their firing per frame (left), then the total duration (middle). At this point, each mean shape is collapsed and plotted over each other. To obtain a subject’s collapse profile, each shape is interpolated to have 100 timepoints and the mean shape is calculated, with SEM based on the each shape’s z-score at each timepoint.

## References

1. Berkes P, Orban G, Lengyel M, & Fiser J (2011) Spontaneous cortical activity reveals hallmarks of an optimal internal model of the environment. Science 331(6013):83–87.

2. Fiser J, Berkes P, Orban G, & Lengyel M (2010) Statistically optimal perception and learning: from behavior to neural representations. Trends in cognitive sciences 14(3):119–130.

3. Cocchi L, Gollo LL, Zalesky A, & Breakspear M (2017) Criticality in the brain: A synthesis of neurobiology, models and cognition. Progress in neurobiology 158:132–152.

4. Beggs JM & Timme N (2012) Being critical of criticality in the brain. Frontiers in physiology 3:163.

5. Palva JM, et al. (2013) Neuronal long-range temporal correlations and avalanche dynamics are correlated with behavioral scaling laws. Proceedings of the National Academy of Sciences of the United States of America 110(9):3585–3590.

6. Hesse J & Gross T (2014) Self-organized criticality as a fundamental property of neural systems. Frontiers in systems neuroscience 8:166.

7. Shriki O & Yellin D (2016) Optimal Information Representation and Criticality in an Adaptive Sensory Recurrent Neuronal Network. PLoS computational biology 12(2):e1004698.

8. Sethna JP, Dahmen KA, & Myers CR (2001) Crackling noise. Nature 410(6825):242–250.

9. Markovic D & Gros C (2013) Power laws and Self-Organized Criticality in Theory and Nature.

10. Virkar Y & Clauset A (2014) Power-law distributions in binned empirical data. The Annals of Applied Statistics 8(1).

11. Tkacik G, et al. (2015) Thermodynamics and signatures of criticality in a network of neurons. Proceedings of the National Academy of Sciences of the United States of America 112(37):11508–11513.

12. Shew WL & Plenz D (2013) The functional benefits of criticality in the cortex. The Neuroscientist : a review journal bringing neurobiology, neurology and psychiatry 19(1):88–100.

13. Marshall N, et al. (2016) Analysis of Power Laws, Shape Collapses, and Neural Complexity: New Techniques and MATLAB Support via the NCC Toolbox. Frontiers in physiology 7:250.

14. Timme NM, et al. (2016) Criticality Maximizes Complexity in Neural Tissue. Frontiers in physiology 7:425.

15. Zimmern V (2020) Why Brain Criticality Is Clinically Relevant: A Scoping Review. Frontiers in neural circuits 14:54.

16. Bak P & Boettcher S (1997) Self-Organized Criticality and Punctuated Equilibria.

17. Beggs JM & Plenz D (2004) Neuronal avalanches are diverse and precise activity patterns that are stable for many hours in cortical slice cultures. The Journal of neuroscience : the official journal of the Society for Neuroscience 24(22):5216–5229.

18. Beggs JM & Plenz D (2003) Neuronal avalanches in neocortical circuits. The Journal of neuroscience : the official journal of the Society for Neuroscience 23(35):11167–11177.

19. Sethna JP (2006) Statistical mechanics : entropy, order parameters, and complexity (Oxford University Press, Oxford ; New York) pp xix, 349 p.

20. Chialvo DR (2010) Emergent complex neural dynamics. Nature Physics 6(10):744–750.

21. de Arcangelis L, Perrone-Capano C, & Herrmann HJ (2006) Self-organized criticality model for brain plasticity. Physical review letters 96(2):028107.

22. Petermann T, et al. (2009) Spontaneous cortical activity in awake monkeys composed of neuronal avalanches. Proceedings of the National Academy of Sciences of the United States of America 106(37):15921–15926.

23. Haimovici A, Tagliazucchi E, Balenzuela P, & Chialvo DR (2013) Brain organization into resting state networks emerges at criticality on a model of the human connectome. Physical review letters 110(17):178101.

24. Stewart CV & Plenz D (2006) Inverted-U profile of dopamine-NMDA-mediated spontaneous avalanche recurrence in superficial layers of rat prefrontal cortex. The Journal of neuroscience : the official journal of the Society for Neuroscience 26(31):8148–8159.

25. Shew WL, Yang H, Petermann T, Roy R, & Plenz D (2009) Neuronal avalanches imply maximum dynamic range in cortical networks at criticality. The Journal of neuroscience : the official journal of the Society for Neuroscience 29(49):15595–15600.

26. Shew WL, Yang H, Yu S, Roy R, & Plenz D (2011) Information capacity and transmission are maximized in balanced cortical networks with neuronal avalanches. The Journal of neuroscience : the official journal of the Society for Neuroscience 31(1):55–63.

27. Gautam SH, Hoang TT, McClanahan K, Grady SK, & Shew WL (2015) Maximizing Sensory Dynamic Range by Tuning the Cortical State to Criticality. PLoS computational biology 11(12):e1004576.

28. Tagliazucchi E, et al. (2016) Large-scale signatures of unconsciousness are consistent with a departure from critical dynamics. *Journal of the Royal Society*, Interface 13(114):20151027.

29. Ma Z, Turrigiano GG, Wessel R, & Hengen KB (2019) Cortical Circuit Dynamics Are Homeostatically Tuned to Criticality In Vivo. Neuron 104(4):655–664 e654.

30. Fagerholm ED, et al. (2015) Cascades and cognitive state: focused attention incurs subcritical dynamics. The Journal of neuroscience : the official journal of the Society for Neuroscience 35(11):4626–4634.

31. Baker FM (1989) Cocaine psychosis. Journal of the National Medical Association 81(9):987, 990, 993-986, passim.

32. Kristensen FW (1994) [Cannabis and psychoses]. Ugeskrift for laeger 156(19):2875–2878, 2881.

33. Stankewicz HA, Richards JR, & Salen P (2022) Alcohol Related Psychosis. *StatPearls*, Treasure Island (FL)).

34. Gardner EL (2011) Addiction and brain reward and antireward pathways. Advances in psychosomatic medicine 30:22–60.

35. Gupta M, Gokarakonda SB, & Attia FN (2022) Withdrawal Syndromes. *StatPearls*, Treasure Island (FL)).

36. Luscher C (2016) The Emergence of a Circuit Model for Addiction. Annual review of neuroscience 39:257–276.

37. Bakhurin KI, Mac V, Golshani P, & Masmanidis SC (2016) Temporal correlations among functionally specialized striatal neural ensembles in reward-conditioned mice. Journal of neurophysiology 115(3):1521–1532.

38. Shobe JL, Claar LD, Parhami S, Bakhurin KI, & Masmanidis SC (2015) Brain activity mapping at multiple scales with silicon microprobes containing 1,024 electrodes. Journal of neurophysiology 114(3):2043–2052.

39. Zare M & Grigolini P (2013) Criticality and avalanches in neural networks. Chaos, Solitons and Fractals: the interdisciplinary journal of Nonlinear Science, and Nonequilibrium and Complex Phenomena 55(Complete):80–94.

40. Schneidman E, Berry MJ, 2nd, Segev R, & Bialek W (2006) Weak pairwise correlations imply strongly correlated network states in a neural population. Nature 440(7087):1007–1012.

41. Dahmen D, Grun S, Diesmann M, & Helias M (2019) Second type of criticality in the brain uncovers rich multiple-neuron dynamics. Proceedings of the National Academy of Sciences of the United States of America 116(26):13051–13060.

42. Cohen MR & Kohn A (2011) Measuring and interpreting neuronal correlations. Nature neuroscience 14(7):811–819.

43. Priesemann V, Munk MH, & Wibral M (2009) Subsampling effects in neuronal avalanche distributions recorded in vivo. BMC neuroscience 10:40.

44. Smith WC, et al. (2016) Frontostriatal Circuit Dynamics Correlate with Cocaine Cue-Evoked Behavioral Arousal during Early Abstinence. eNeuro 3(3).

45. Friedman N, et al. (2012) Universal critical dynamics in high resolution neuronal avalanche data. Physical review letters 108(20):208102.

46. Chen BT, et al. (2013) Rescuing cocaine-induced prefrontal cortex hypoactivity prevents compulsive cocaine seeking. Nature 496(7445):359–362.

47. Flagel SB, Akil H, & Robinson TE (2009) Individual differences in the attribution of incentive salience to reward-related cues: Implications for addiction. Neuropharmacology 56 Suppl 1:139–148.

48. Zapperi S, Baekgaard Lauritsen K, & Stanley HE (1995) Self-organized branching processes: Mean-field theory for avalanches. Physical review letters 75(22):4071–4074.

49. Landmann S, Baumgarten L, & Bornholdt S (2021) Self-organized criticality in neural networks from activity-based rewiring. Physical review. E 103(3-1):032304.

50. Nigam S, et al. (2016) Rich-Club Organization in Effective Connectivity among Cortical Neurons. The Journal of neuroscience : the official journal of the Society for Neuroscience 36(3):670–684.

51. Reed WJ & Hughes BD (2002) From gene families and genera to incomes and internet file sizes: why power laws are so common in nature. Physical review. E, Statistical, nonlinear, and soft matter physics 66 6 Pt 2:067103.

52. Sheynin OB (1979) C.F. Gauss and the theory of errors. Archive for History of Exact Sciences 20(1):21–72.

53. Ahern J, Stuber J, & Galea S (2007) Stigma, discrimination and the health of illicit drug users. Drug and alcohol dependence 88(2-3):188–196.

54. Mayer AR, Wilcox CE, Teshiba TM, Ling JM, & Yang Z (2013) Hyperactivation of the cognitive control network in cocaine use disorders during a multisensory Stroop task. Drug and alcohol dependence 133(1):235–241.

55. Jasinska AJ, Chen BT, Bonci A, & Stein EA (2015) Dorsal medial prefrontal cortex (MPFC) circuitry in rodent models of cocaine use: implications for drug addiction therapies. Addiction biology 20(2):215–226.

56. Hagen E, et al. (2015) ViSAPy: a Python tool for biophysics-based generation of virtual spiking activity for evaluation of spike-sorting algorithms. Journal of neuroscience methods 245:182–204.

57. Tlaie A, Shapcott KA, Tiesinga P, Schölvinck ML, & Havenith MN (2021) Does the brain care about averages? A simple test. bioRxiv:2021.2011.2028.469673.

58. Tononi G, Sporns O, & Edelman GM (1994) A measure for brain complexity: relating functional segregation and integration in the nervous system. Proceedings of the National Academy of Sciences of the United States of America 91(11):5033–5037.

59. Bonmati E, Bardera A, Feixas M, & Boada I (2018) Novel Brain Complexity Measures Based on Information Theory. Entropy 20(7).

60. Priesemann V, et al. (2014) Spike avalanches in vivo suggest a driven, slightly subcritical brain state. Frontiers in systems neuroscience 8:108.

61. di Santo S, Villegas P, Burioni R, & Munoz MA (2018) Landau-Ginzburg theory of cortex dynamics: Scale-free avalanches emerge at the edge of synchronization. Proceedings of the National Academy of Sciences of the United States of America 115(7):E1356–E1365.

62. Ma YY, et al. (2014) Bidirectional modulation of incubation of cocaine craving by silent synapse-based remodeling of prefrontal cortex to accumbens projections. Neuron 83(6):1453–1467.

63. Tannenbaum J & Bennett BT (2015) Russell and Burch’s 3Rs then and now: the need for clarity in definition and purpose. Journal of the American Association for Laboratory Animal Science : JAALAS 54(2):120–132.

64. Lansey JC (2022) Power Law, Exponential, and Logarithmic Fit. MATLAB Central File Exchange.

